# pyKinaXe: a fast and robust turnkey kinase activity profiler

**DOI:** 10.64898/2026.05.12.724658

**Authors:** David Wuttke, Eberhard Hildt, Pavel V. Kolesnichenko

## Abstract

Peptide microarray technologies such as PamDx enable direct measurement of peptide phosphorylation by upstream kinases, yet extraction of kinases from raw data depends on proprietary software or separate open-source alternatives delivering time-consuming processing across a variety of different steps, limiting throughput for experimental large-scale kinome generation in clinical and research settings and compound/drug screening. We developed pyKinaXe, a Python package for automated end-to-end analysis of PamChip^®^ data, integrating robust image processing, kinase analysis, and pathway enrichment into a single one-click pipeline. We validated pyKinaXe on two independent datasets against existing peptide-microarray-based kinase activity inference tools, demonstrating qualitative agreement and, at the same time, drastic speed-up (6-fold for image processing and 42-fold for statistical kinase inference). pyKinaXe addresses the key limitations of existing tools (slow inference, fragmented workflows, and poor usability), enabling fast and robust analysis, and facilitating high-throughput experiments and large-scale kinome profiling. pyKinaXe is implemented in Python 3.13 and distributed under the Apache 2.0 License. Source code, documentation, and installation instructions are freely available at https://github.com/pykinaxe/pyKinaXe. The benchmark data are available at Mendeley Data (doi: 10.17632/ynp7f92n47.1). pyKinaXe’s user-friendly web-based interface can be accessed at https://pykinaxe.github.io/home.

## Introduction

Protein kinases regulate target proteins via phosphorylation, thus controlling a broad variety of cellular processes [33]. They have become high-priority therapeutic targets, with more than 115 kinase inhibitors already approved for clinical use [73; 54; 72]. Most approved inhibitors target neoplastic diseases, where the severity of the disease may justify accepting greater treatment-related risks than in non-oncological indications. Consequently, the clinical use of kinase inhibitors outside oncology remains limited.

Identifying active kinases for a particular biological context of interest remains a challenge due to the complexity of signaling networks and the dynamic nature of phosphorylation [56]. Classical approaches such as Western blotting [80] use phospho-specific antibodies to detect kinase phosphorylation at activation sites as a surrogate marker for kinase activity [50]. Although straightforward and widely established, this strategy is limited to a small number of pre-selected kinases per experiment, and depends on the availability and specificity of phospho-antibodies reflecting kinase activity [2; 7]. Mass spectrometry (MS)-based methods can quantify tens of thousands of phosphorylation sites [10;32; 9], resulting in large phosphoproteomic datasets. However, they can measure phosphorylated proteins but not direct kinase activity and therefore require computational inference and a very sophisticated infrastructure to link phosphorylation signatures to the corresponding kinases [66; 25]. In contrast, peptide microarray technologies such as PamDx (PamGene International B.V., formerly marketed as PamGene), allow direct measurement of kinase activity using immobilized substrate peptides, combining small sample requirements (1–5 *µ*g per microarray), real-time acquisition of phosphorylation kinetics, and the capacity for high-throughput experimentation [6; 3; 60; 61]. Despite these advantages, data from peptide microarrays require careful image processing and non-trivial statistical deconvolution [6; 3], and the time-consuming nature of current analysis workflows limits the high-throughput capability that the technology itself offers.

The past 25 years have witnessed the rapid evolution of databases, computational methods, and benchmarking for kinase inference [28; 75; 66; 23; 55], mostly targeting data obtained by means other than PamDx kinase activity profiling. The first generation of computational methods [57; 92; 31; 47] focused on predicting kinase–substrate relationships (KSR) solely on the basis of protein sequence logic, ultimately generating KSR databases. These in turn triggered the next generation, which is by far the largest, most active and most heterogeneous group of methods [14; 87; 53; 88; 39;26; 90; 24; 82; 49; 5] aimed at predicting kinase activity from MS-derived phosphoproteomic data. This generation pioneered statistical approaches applied to KSR databases, such as KSEA [14] and PTM-SEA [39], and later introduced database-agnostic solutions [82; 49]. The third generation of methods [16; 40; 51; 91; 12] such as KRSA [20] and KEA3 [41] operates on non-phosphoproteomic data (gene lists, transcriptome, multiome, drug compounds, peptide microarrays) and is increasingly dominated by machine learning approaches.

Peptide microarray data generated with PamDx technology occupy a unique position among the data used by third-generation methods. The peptide microarrays are provided on a chip (PamChip^®^) and contain selected peptide sequences placed inside porous membranes, allowing cell lysates to be actively cycled through while fluorescently labeled phosphoSer/phosphoThr or phosphoTyr-specific antibodies detect phosphorylation in real-time [6; 3]. The technology supports two different PamChip^®^ microarrays, with substrate peptides specific to tyrosine kinases (PTK) [89] and those specific to serine/threonine kinases (STK) [35]. The raw data are images of PTK/STK microarrays acquired by a CCD camera capturing fluorescence differences between control and test conditions. The analysis of such data has remained largely dependent on proprietary BioNavigator^®^ software [59]. Although the software provides complete data analysis workflows (including both image and statistical analysis), the processing is time consuming, and multiple manual adjustments are required at different processing stages. Together with its limited accessibility, this impedes extensibility and transparency, creating a bottleneck for research, clinical translation, and high-throughput analyses. To address these limitations, only two open-source solutions have emerged so far: pamgeneAnalyzeR[8] and KRSA [20], both implemented in R. The pamgeneAnalyzeR package, however, focuses exclusively on image analysis of peptide microarray data and does not provide subsequent statistical analysis. Similarly, KRSA lacks integrated image processing and requires pre-quantified data to perform statistical analysis. Other tools, such as the Kinomics Toolbox [22], start from BioNavigator^®^ exports and provide pre-processing and differential peptide analysis, but neither image analysis nor kinase analysis. To date, no tool performs both image and statistical analysis in a single, uninterrupted pipeline.

Here we describe the development and validation of pyKinaXe, a novel tool for end-to-end analysis of PamChip^®^ peptide microarray data. pyKinaXe occupies a unique niche in the landscape of kinase inference toolkits by providing accurate, fast, and fully automated one-click analysis from raw fluorescence images to interpretable kinase activity profiles. In the complete analytical pipeline, pyKinaXe integrates statistically robust image processing strategies, quantification of phosphorylation kinetics, multi-database substrate-kinase mapping, and explainable statistics-based kinase analysis for both kinase families and individual kinases. To further facilitate accessibility, we developed a turnkey web interface for pyKinaXe, available at https://pykinaxe.github.io/home.

## Materials and methods

### Required input

The only input required by pyKinaXe is that which is automatically generated during the experiment (Figure 1). In addition to raw images, this includes two text files describing layouts of the peptide microarrays and sample annotations. The user carrying out a PamDx experiment only needs to follow a sample annotation convention ([c|t]#_SampleName_<bio>_<tech>, where SampleName refers to the sample’s meaningful name, <bio>/<tech> corresponds to its biological/technical replicate number, and [c|t]# explicitly indicates whether it is a Control or Test sample; in [c|t]#, the number can be higher than 4 to indicate the case in which more than 4 conditions were used across the chips in an experiment) for pyKinaXe to correctly parse the sample annotation file. The experimentally generated data can also be provided to pyKinaXe via the Internet and analyzed online.

**Figure 1.**
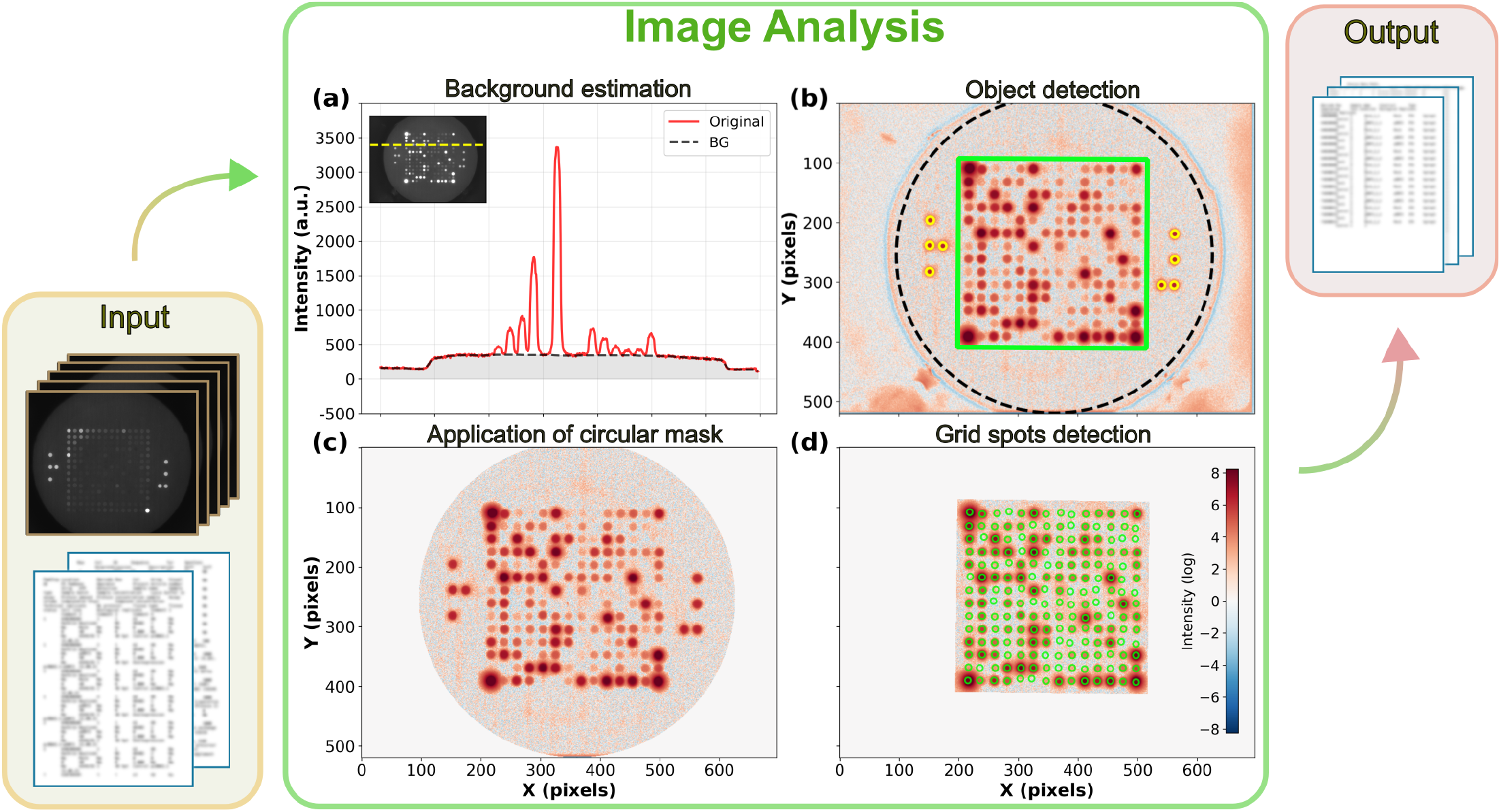
Image analysis pipeline with inputs (raw images and two text files) and outputs (text and tabulated files) indicated. (a) A cross-section of a raw image (inset) taken along the indicated line (yellow dashes), showing the estimated BG (black dashes) beneath the signal curve (solid red). (b) Detection of the reference spots (yellow circles) and the main microarray grid (green square). (c) Detection of the optical aperture and application of the corresponding circular mask (black dashes). (d) Detection of the main microarray spots (green circles) demonstrated on an image with the square mask from (b) applied. Background-subtracted images are used in (b–d), where intensity values are symmetrically log-transformed. The colorbar in (d) applies equally to (b) and (c).

### Generation of image data

Each PamChip^®^ contains four microarrays, each holding a grid of peptide spots. The spots contain immobilized peptides and are located on a porous 3D membrane [60; 61]. During the experiment, cell lysate is repeatedly pumped through the membranes, allowing kinases in the sample to phosphorylate the peptides while fluorescently labeled antibodies detect the phosphorylation signal in real time. At each pump cycle, microarray images are captured at multiple exposure times to ensure that high- and low-intensity signals are accurately recorded, thereby increasing the dynamic range of measurement [70]. The raw-image capture protocol is split into pre-wash and post-wash phases. Pre-wash images are acquired during those pumping cycles when kinase reactions and antibody binding take place, building up phosphorylation signal over time. Post-wash images are acquired after unbound antibodies have been removed (washed out), providing a cleaner readout of phosphorylation without non-specific contributions [60; 61].

### Image processing

#### Correction for uneven background

Prior to the analysis of fluorescence from antibodies, the background (BG) signal, arising, *e*.*g*., from non-specific fluorescence, dark current, or stray light, must be accounted for. Unlike pamgeneAnalyzeR (which estimates BG at each microarray spot from fixed locations within the optical aperture (OA)) and BioNavigator^®^ (that uses raw signal around grid spots as a BG estimate), pyKinaXe can account for the slowly varying, spatially uneven signal distribution across the entire field of view (Figure 1a) via a three-step morphological grey opening (MGO) [27], as implemented in OpenCV [11], applied to Gaussian pre-smoothed [85] images to exclude noise from such estimation. Each consecutive step is characterized by a larger structural element used in MGO, ensuring that no residual fluorescence contributes to the estimated BG. pyKinaXe additionally accounts for “cross-talk” cases in which bright saturated spots add additional signal to the surrounding weak spots. This is achieved by estimating later (at a stage when grid spots have already been identified) the raw signal around each grid point and averaging it with the MGO estimated baseline (see Supplementary Material for more details).

#### Identification of objects in images

Each image contains distinct objects to be identified: the reference (REF-)spots, the main microarray grid, and the optical aperture (Figure 1b).

REF-spots are located to the left and right of the main microarray grid, arranged in patterns resembling “TeeWee” (T-shape, left) and “Blue Ricky” (J-shape, right). These REF-spots emit bright fluorescence, ensuring that even when signals from the main microarray grid are weak, the grid can still be unambiguously localized by the algorithm. Accurate and statistically robust identification of the REF-spot locations is therefore a critical step in the pipeline.

We detect the REF-spots in two stages. First, local intensity maxima are identified by applying a maximum filter to Gaussian-smoothed images, retaining only those peaks that are above the 80th percentile of intensity. Second, the candidates in the left and right halves of the image are considered, and a global search is performed: for each candidate in the left (right) half, the algorithm attempts to verify a T-shaped (J-shaped) pattern using *a priori* knowledge of geometrical relations between the four spots within each pattern. Potential T–J pairs are then validated against their expected relative positions. If at least one of the REF-spots is not identified (e.g., due to an excess of dust), the algorithm is tried on a median-filtered image.

Once the REF-spots are localized, the position of the main microarray grid is determined from geometric considerations (Figure 1b), accounting for its rotation angle arising from the finite angular tolerance of the PamChip^®^ within its slot on the measuring device.

The next distinct object to be identified is OA (Figure 1c), which encapsulates both the REF- and microarray spots. Because the position of OA varies between experiments, we detect OA from the image content, using contrast-enhanced MGO-processed images (featuring plateaus without spots). Otsu’s method [58] is then applied for automatic thresholding: the initial center is estimated from the centroid of the resulting image, and the area of OA is estimated from the largest connected region. The circle is then refined: 180 rays (corresponding to 180 degrees) are thrown from the centroid, and bright-to-dark transitions are detected. With outliers rejected, the detected transition points are fit with a circle, resulting in robust estimates of OA centers.

A circular mask of slightly reduced radius (to reduce the OA boundary’s edge effects) is then applied to all images, setting signals outside the OA to zero.

#### Detection of the microarray spots

Finally, the individual microarray grid spots (Figure 1d) are identified in several stages. First, theoretical spot locations are assumed using the estimated rotation angle of the grid and *a priori* knowledge of inter-spot spacing. Next, global grid refinement is performed using an optimizer (L-BFGS-B [93] and Powell [67] as implemented in SciPy [85]) in which global parameters such as overall grid position, its rotation angle, and inter-spot spacing are iteratively varied to maximize total integrated fluorescence intensity from all spots. Finally, to account for the spots that deviate from their assumed positions, a second optimization of individual spot positions within the grid is carried out. Having identified the location of the grid spots, the median intensity of each spot is derived for the subsequent kinase inference.

### Kinase inference

#### Identification of the experimental setup

Prior to the statistical analysis, control and test samples on each chip are automatically identified. pyKinaXe supports experimental setups with multiple control and multiple test conditions and, by default, forms every control-versus-test pair. STK and PTK data are analyzed separately due to differences in their experimental design and signal levels. Accordingly, the QC protocols differ between the two microarray types, while the downstream peptide statistics and kinase enrichment analysis follow identical workflows (Figure 2).

**Figure 2.**
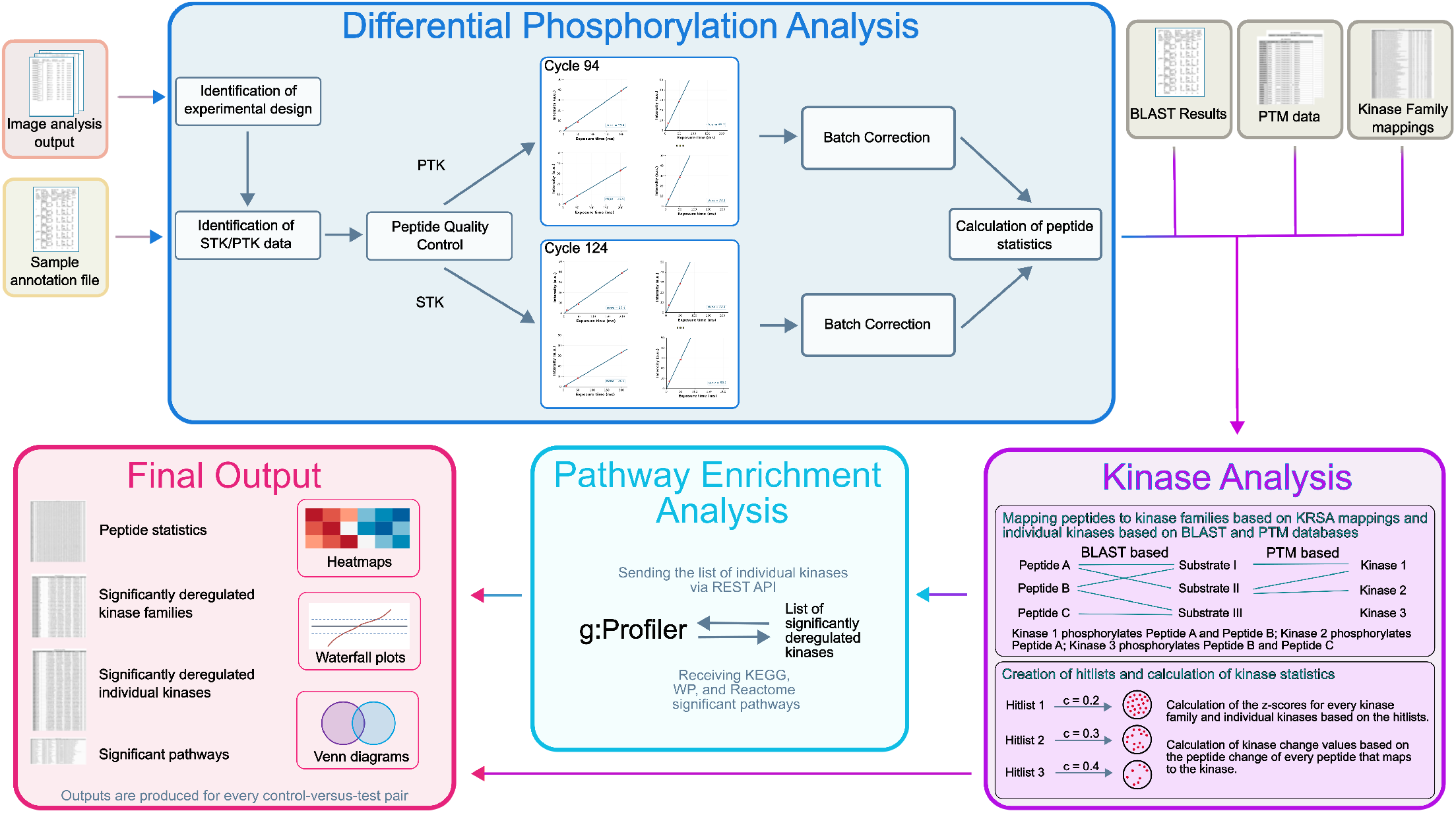
Workflow of the statistical analysis with inputs (sample annotation file and the image analysis output) and outputs (peptide level, kinase family level, individual kinase level, and pathway level) indicated. Differential phosphorylation analysis, kinase analysis, and pathway enrichment analysis are shown as the building blocks of the statistical analysis pipeline.

#### Quantifying phosphorylation intensity of peptides

Within each pump cycle, peptide phosphorylation is captured at several exposure times. Similarly to BioNavigator^®^ [62], for each pump cycle we quantify peptide phosphorylation magnitude by extracting the slope of linear dependence of median fluorescence intensity *I* on exposure time *T*, as follows:

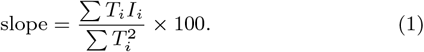

An additional multiplication by 100 ensures numerical stability in further analysis. The absence of a linear intercept is expected (no signal at *T* = 0). The calculation of the slope yields a single measure of phosphorylation per peptide per cycle.

#### Quality control

Prior to slope calculation, measurements with signal saturation levels of *≥*5% are excluded to increase the dynamic range of integrated signals [70]. After this, by default, a linear regression of spot signal against exposure time is fitted for every spot at the end-point cycle. A peptide is retained only if the coefficient of determination of that regression is *≥*0.9. These criteria closely reproduce the filter of KRSA [20], so that the retained peptide set is comparable to the one KRSA operates on.

Alternatively, a quality control method close to BioNavigator^®^’s approach can be applied.

#### Differential phosphorylation analysis

After QC, PTK- and STK-peptide slopes are log_2_-transformed for downstream analysis. This transformation places up- and down-regulated signals on a symmetric scale [69]. For each peptide *g*, differential phosphorylation between test and control is calculated as peptide change Δ_*g*_, defined as the difference between the mean log_2_-transformed slopes of the test and control group:

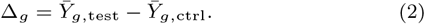

Because samples are distributed over several PamChips and each chip carries its own additive offset, pyKinaXe regresses out a per-chip effect before forming the contrast, so that Δ_*g*_ is evaluated on batch-adjusted values.

#### Peptide-kinase mapping

In order to map the peptides to the individual kinases, two steps are required. First, peptides are linked to the human substrate proteins they represent using the Basic Local Alignment Search Tool (BLAST) [4; 13], retaining only matches above a user-configurable sequence similarity threshold (by default 80%). Second, substrates are linked to phosphorylating kinases using UniProt [77], iPTMnet [30], and OmniPath [81] databases that contain information on post-translational modifications (PTM) of proteins (including PhosphoSitePlus [29], SIGNOR [48], PhosphoELM [21], HPRD [65], and dbPTM [19]). A kinase is linked to a peptide if any of its BLAST-matched proteins appears as a substrate of that kinase in the PTM database. Analysis of phosphatases is not included in the peptide-kinase mapping, as phosphatase inhibitors are included in the sample preparation protocol for PamChip^®^ microarrays [60; 61].

To map the peptides on the array to the kinase families, a table from KRSA is used [20]. This table is highly detailed and allows for a comparison of the results of pyKinaXe with those of KRSA.

#### Kinase–peptide enrichment analysis

In pyKinaXe, kinase activity is inferred using a hypergeometric kinase–peptide enrichment analysis (KPEA) algorithm inspired by KRSA [20]. The KPEA is conducted on two layers, first for kinase families (based on KRSA), and second for individual kinases. In contrast to KRSA, which reports only kinase families, pyKinaXe additionally resolves kinase–peptide relationships at the level of individual kinases, thus enabling a higher kinase resolution. KPEA is performed on a subset of QC-passed peptides that can be mapped to at least one kinase (or, in the case of the kinase family analysis, to at least one kinase family). Hit lists are then generated based on Δ_*g*_ and a cutoff *c*,

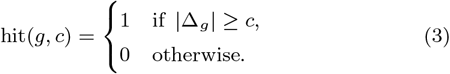

By default, cutoffs *c* are set to 0.2, 0.3, and 0.4, matching the default values used in KRSA [20]. For each kinase (or kinase family) *k* and cutoff *c*, the number of peptides that map to kinase (or kinase family) *k* and appear in the observed hit list is defined as 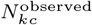. A null distribution is constructed by drawing a random peptide set of the same size as the observed hit list, without replacement, from the background pool of peptides. By default, the background pool is the set of peptides used on the respective PamChip that are present in the kinase mapping reference. Restricting the pool to the QC-passed peptides as a background is available as an alternative option. Lastly, the observed count of peptides that map to a kinase or kinase family within each hitlist is standardized against the null distribution. The *z*-score for kinase *k* at cutoff *c* is calculated as

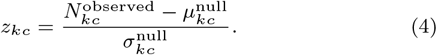

The final score is obtained by averaging the per-cutoff *z*-scores across the hit lists. By default, kinase families are considered significant when the averaged score deviates from the null distribution by more than 2.0 standard deviations (i.e., when 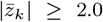). The corresponding threshold in the analysis of individual kinases is 1.3, reflecting the exploratory character of that layer.

A positive *z*-score indicates that kinase *k* is represented by a larger number of hit peptides than expected by chance, whereas a negative *z*-score indicates under-representation. Additionally, kinase change is calculated as the mean of the peptide change values across all peptides assigned to a given kinase to quantify the overall shift in kinase activity between conditions. PTK and STK arrays are scored independently and reported separately for each control-versus-test pair. For the downstream pathway enrichment, PTK and STK results of each comparison are merged.

#### Pathway enrichment analysis

Kinase gene symbols are submitted to g:Profiler [38] via API, querying KEGG [36], WikiPathways [1] and Reactome [52] for human pathways that contain these kinases. Pathway enrichment is performed using g:Profiler/g:GOSt (version e114 eg62 p19 27110d83) with a g:SCS-corrected threshold of 0.05. By default, the background in the enrichment is set to the g:Profiler web values. Additionally, the background can be set to include only kinases that can be mapped using our peptide-kinase mapping.

### Final output

pyKinaXe produces output for both the image analysis and the KPEA. For image analysis, CSV files are generated, separated by PTK and STK, which contain the intensity levels of the spots organized by array, spot, exposure time, and cycle. This output is comparable to the output from BioNavigator^®^’s image analysis.

For the KPEA, results are generated at the peptide level, pathway level, kinase level, and kinase family level for each control-versus-test pair. Additionally, at peptide level, pyKinaXe produces heatmaps showing the log_2_-transformed slopes of every peptide on the corresponding samples and waterfall plots of the log2 fold changes of every peptide. Also, on pathway level, heatmaps are created showing the kinase change of significantly deregulated kinases for all significantly affected pathways.

### Optimization for algorithm’s speed

To ensure computational efficiency, we applied several optimization strategies throughout the algorithm, including just-in-time (JIT) compilation via Numba [43], parallelization of independent tasks using Joblib [83], image downsampling prior to computationally intensive operations, batch processing to minimize overhead, and vectorization of mathematical operations.

## Results

To evaluate pyKinaXe, we re-analyzed the kinome [79] published by Thiyagarajah et al. [78], extracted via BioNavigator^®^ from the human hepatoma cell line Huh7 cells ectopically expressing large hepatitis delta antigen (LHDAg), small hepatitis delta antigen (SHDAg), or the replicating hepatitis delta virus’ (HDV) genome via the complementary DNA (cDNA) clone of pSVLD3 (HDV cDNA expression plasmid) [42]. Each condition was measured on two PamChips, i.e. n = 2 technical replicates per condition on both the PTK and STK arrays.

### Validating image analysis against BioNavigator

To validate the image analysis of pyKinaXe, we compared the spot intensities produced by the image processing analysis against the output of BioNavigator^®^’s image analysis (Figure 3). For both PTK (a) and STK (b), the spot intensities behave similarly to those from BioNavigator^®^, with a small systematic offset. pyKinaXe reports slightly higher values at low intensities, mainly because of its averaged background estimate. A more detailed comparison of the output of pyKinaXe against BioNavigator^®^ can be found in the Supplementary Material.

**Figure 3.**
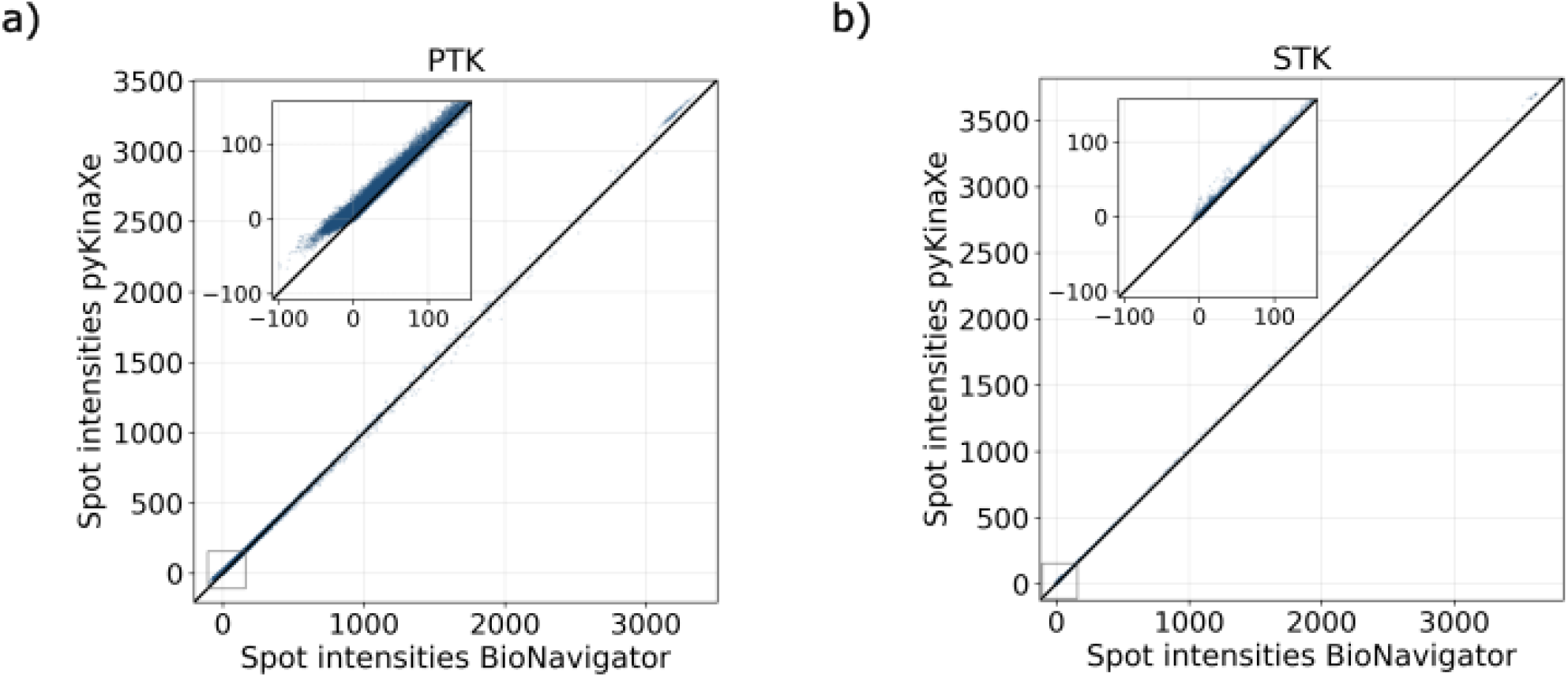
Comparison of spot intensities extracted by pyKinaXe (y-axis) and BioNavigator^®^ (x-axis), based on data from Thiyagarajah et al. [78]. A magnified view shows the spot intensities at lower intensities. (a) shows the spot intensities for PTK chips, and (b) shows the intensities for STK chips.

### Benchmarking on experimental results

To evaluate the KPEA from pyKinaXe, we compared the results at the kinase family level with the results from KRSA. For this purpose, KRSA was run on the image analysis output from pyKinaXe. The results of this analysis are shown in Figure 4. The waterfall plot shows the differences between the results from pyKinaXe and KRSA for each kinase family. The results show very similar *z*-scores for the kinase families. In addition, most of the kinase families identified as significantly deregulated by KRSA are also identified by pyKinaXe (23 out of 30). For LHDAg, KRSA and pyKinaXe show substantial agreement, with 10 kinase families identified by both approaches. In addition, KRSA detects 4 kinase families that are not captured by pyKinaXe, whereas pyKinaXe identifies one kinase family exclusively. For SHDAg, 2 kinase families are identified by both KRSA and pyKinaXe. Additionally, 2 kinase families are detected only by KRSA. For HDV, a comparison reveals that 11 kinase families are identified by both approaches, while one kinase family is detected exclusively by KRSA and one kinase family exclusively by pyKinaXe. In addition to the analysis presented here, the Supplementary Material includes another analysis based on a second publicly available dataset [34; 44].

**Figure 4.**
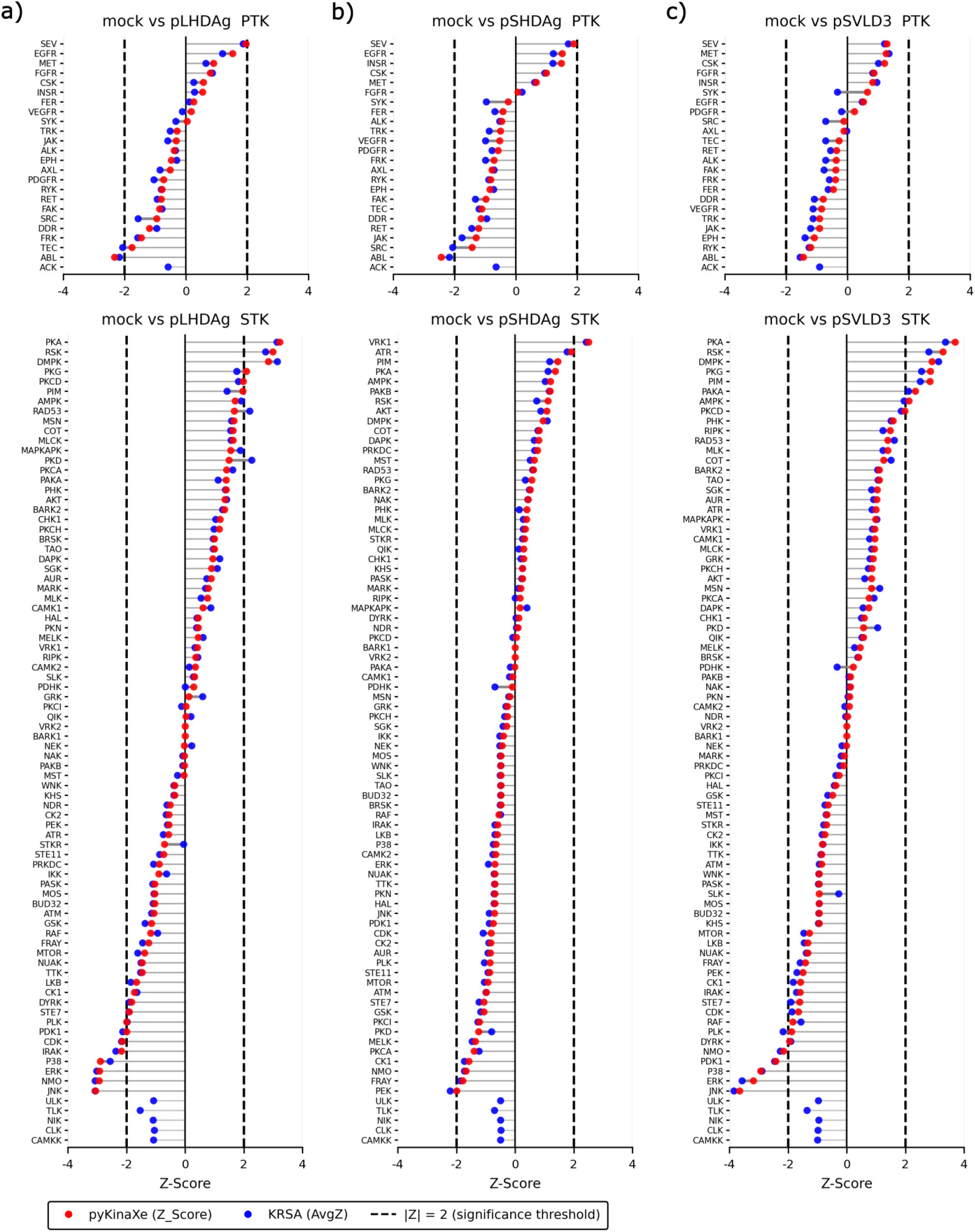
Comparison of the kinase family *z*-scores between pyKinaXe and KRSA based on the pyKinaXe image analysis output of the data from Thiyagarajah et al. [78]. The x-axis shows the *z*-score, and the y-axis shows the various kinase families (sorted by *z*-score). (a) shows waterfall plots comparing mock to pLHDAg for both PTK and STK chips. (b) shows the results for pSHDAg and (c) for HDV. The blue points represent the *z*-scores from KRSA, and the red points represent the *z*-scores from pyKinaXe. The significance levels (dashed vertical lines) are *−*2 and +2.

On the individual kinase level, pyKinaXe yields 49 significant kinases for LHDAg (10 PTK, 39 STK), 53 for HDV (13 PTK, 40 STK) and 11 for SHDAg (7 PTK, 4 STK), for a union of 76 distinct kinases across the three control-versus-test pairs (Figure 5a). LHDAg and HDV share 30 of their significant kinases, whereas SHDAg overlaps with either of them only marginally (5 and 6 kinases). Four kinases (FLT1, PBK, PKN2 and PRKACA) are identified as significant in all three control-versus-test pairs, while 18, 21 and 4 kinases are specific to LHDAg, HDV and SHDAg, respectively. Mapping the significant kinases onto KEGG pathways returned 82 enriched pathways for LHDAg (Figure 5b), 103 pathways for HDV (Figure 5c), and a single pathway for SHDAg (Figure 5d). MAPK signaling is therefore the only pathway enriched in all three control-versus-test pairs, while LHDAg and HDV share 68 pathways.

**Figure 5.**
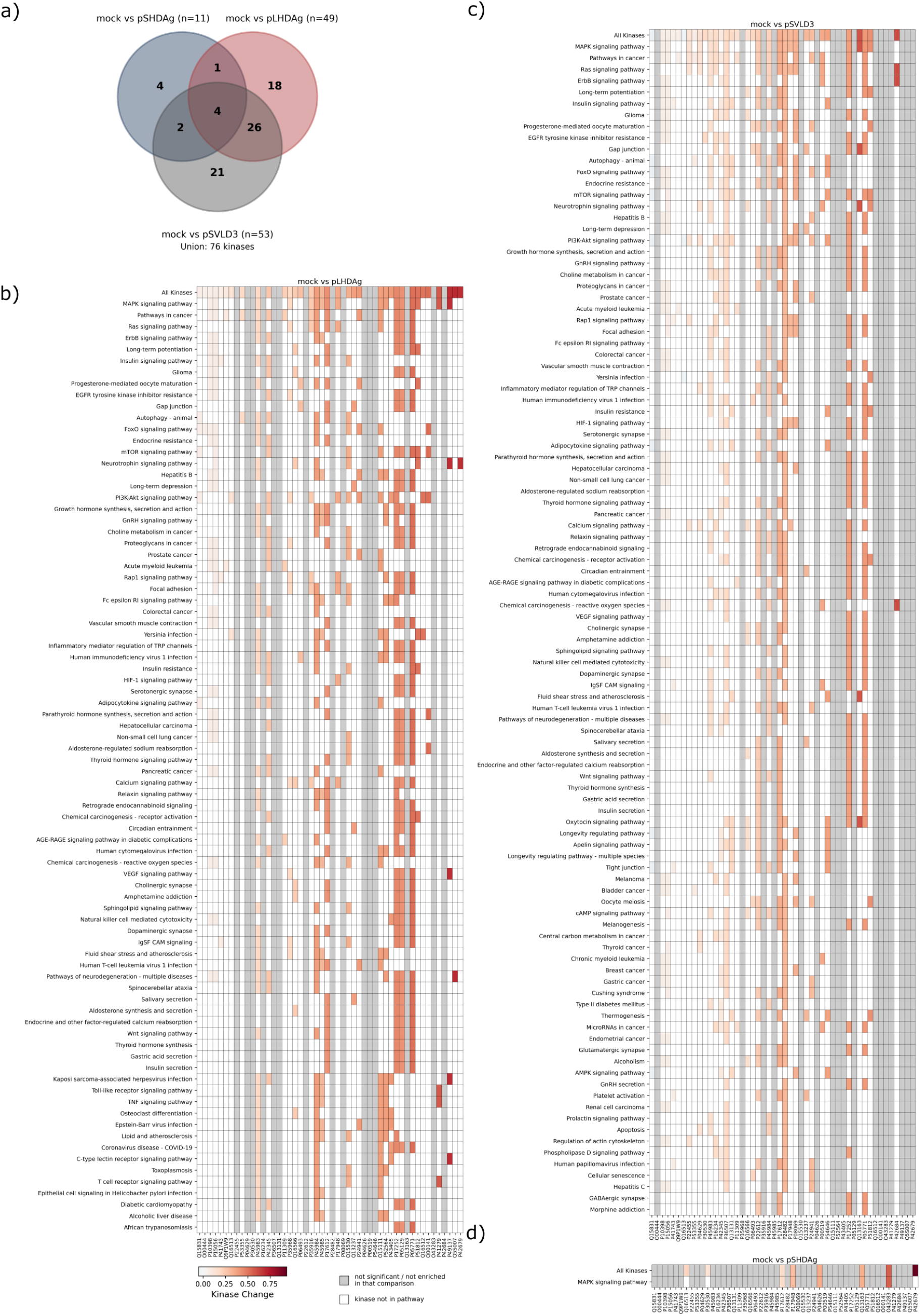
pyKinaXe benchmarking results at the individual kinase and pathway levels. (a) shows the overlap of individual kinases between the three control-versus-test pairs (LHDAg, SHDAg and HDV) in the form of a Venn diagram. (b) shows a heatmap presenting the significantly affected pathways of mock versus LHDAg, together with the significantly identified kinases in these pathways. The heat scale shows the kinase change. (c) shows the pathway results for HDV and (d) for SHDAg.

### Runtime efficiency and robustness

The total runtime for a representative experimental dataset (560 images, 697*×*520 pixels) is about 24 seconds on a laptop with a 10-core Apple M4 processor, with image analysis and statistical analysis accounting for 20 and 4 seconds, respectively. We compared pyKinaXe’s efficiency with that of the published open-source alternatives, pamgeneAnalyzeR for image analysis and KRSA for kinase analysis (both published in R), run on the same machine and the same input images. With pamgeneAnalyzeR run using its default settings (running on STK only), and KRSA configured as in its own report template (running on BioNavigator^®^ input), pamgeneAnalyzeR required 128 seconds and KRSA 170 seconds, making pyKinaXe (at least) 6 times faster at image analysis and 42 times faster at the statistical analysis on the tested input data.

Additional tests on the robustness of the pyKinaXe image analysis can be found in the Supplementary Material.

### User interface

The user interface allows users to upload both PTK and STK raw data. With a single click, the analysis can be performed. Once the analysis is complete, the interface shows lists of significantly deregulated kinases and peptide heatmaps, while the complete set of all results can be downloaded.

## Discussion

pyKinaXe provides the first open-source tool that integrates image processing and statistical analysis for kinase inference into a single uninterrupted (i.e., without manual adjustment steps) speed-optimized pipeline for PamChip^®^ peptide microarray data. An additional web-based user interface further enhances accessibility of peptide-microarray-based kinase activity profiling to a wider range of users. Thus, pyKinaXe addresses the main limitations of existing solutions that hinder widespread adoption, namely long inference times, lack of streamlined workflows, and a steep usability barrier, enabling its use in clinical and research settings as well as in high-throughput screening.

Validation of the reported HDV data shows that pyKinaXe produces similar results in both image analysis and kinase family analyses. The image analysis results resemble those of BioNavigator^®^ closely, with only a slight difference (Figure 3). This offset can be explained by the differences in estimating median fluorescence intensities from each of the grid spots as well as background signal contributions. The kinase family analysis produces results that show only minor discrepancies from those generated using KRSA (Figure 4 and Supplementary Material), which further validates the correct implementation of our tool. Again, small differences in the results can be explained by the different methodological approaches of KRSA compared to pyKinaXe. The benchmarking results show that pyKinaXe cannot generate a score for one kinase family in one case for PTK data, and for five kinase families in the STK data. In our approach, these families are removed during quality control, because their substrate peptides either fall below the signal thresholdor have coefficients of determination of the linear regression *<* 0.9. Validation on an additional, independent dataset showed qualitatively similar PTK and STK spot-intensity patterns between pyKinaXe and BioNavigator^®^, as well as consistent kinase family *z*-scores between pyKinaXe and KRSA (Supplementary Material Figures 6 and 7).

For the validation of the statistical analysis, we compared the output at the kinase family level by pyKinaXe with the results from KRSA (on pyKinaXe image analysis). The results show a high degree of agreement between the *z*-scores generated by both approaches, with kinase families identified as significant by pyKinaXe being widely supported by the literature. Members of the JNK and ERK kinase families, which were identified in both LHDAg and HDV, have previously been implicated in hepatitis-related processes or HDV replication. In particular, JNK signaling has been associated with liver disease [76], while ERK1/2, members of the ERK family, have been shown to promote HDV replication [17]. Furthermore, CDK8 and CDK19, members of the CDK family identified in LHDAg, are required for HDV replication [68]. Members of kinase families identified as significant in both LHDAg and HDV have also been implicated in hepatocellular carcinoma (HCC). For example, JNK1, a member of the JNK family, has been associated with hepatocarcinogenesis [74], while RSK2, a member of the RSK family, is recurrently mutated in HCC [15]. Members of kinase families identified by pyKinaXe exclusively in HDV have also been associated with HCC. For example, PIM1 and PAK1, members of the identified PIM and PAKA kinase families, respectively, have been implicated in HCC [46; 18]. Additionally, members of kinase families identified in SHDAg have been associated with HCC, such as VRK1 [45].

The final output scores at the individual kinase level of pyKinaXe and BioNavigator^®^ are not interchangeable, because pyKinaXe deliberately reports a different and more directly interpretable quantity. BioNavigator^®^’s ‘Final Score’ is the sum of two permutation probabilities, the Specificity Score (permutation of peptides) and the Significance Score (permutations of samples) [64]. pyKinaXe instead reports for each kinase (or kinase family) the number of its substrate peptides among the peptides whose Δ_*g*_ exceeds a cutoff (0.2, 0.3, 0.4; averaged), standardized against the mean and standard deviation of the null defined by the peptides printed on the PamChips. Thereby, pyKinaXe quantifies the extent to which a kinase’s (or kinase family’s) substrate set is over- or under-represented among the most strongly changed peptides, relative to the entire peptide population on the chip, thereby nominating kinases whose substrate sets could account for the observed phosphorylation pattern. The *z*-scores from pyKinaXe and the ‘Final Score’ from BioNavigator^®^ therefore measure different things on different scales.

Nevertheless, pyKinaXe can identify a broad range of individual kinases that have been linked to HDV or HCC in the literature. CDK19 [68] and MAPK1 [17] (both identified in HDV) are related to HDV replication. Individual kinases like RPS6KA3 [15] and MTOR [84] (both identified in LHDAg and HDV), and FGFR4 [86] (identified in HDV) are related to HCC. At the pathway level (based on individual kinases), pyKinaXe also identifies signaling pathways that are consistent with the literature. Thus, PI3K-Akt signaling (identified in LHDAg and HDV) and MAPK signaling (identified in LHDAg, SHDAg, and HDV) are recognized as significant, which is consistent with the original study of the input data [78].

PamDx’s UKA algorithm assigns individual kinases to substrates using six PTM databases [71; 63], including PhosphoNET for in silico-predicted interactions based on kinase sequences [37]. These predictions nominate candidate kinases but do not establish that they act as upstream regulators in the biological context analyzed. Five of the databases used by PamDx are also integrated into the data sources used by pyKinaXe for the individual kinase analysis. However, PhosphoNET is not openly accessible for download and is therefore not integrated into pyKinaXe as an open-source tool. This difference in the underlying kinase–substrate interactions likely leads to deviations at the level of individual kinases. Alongside the PTM databases used by PamDx, pyKinaXe includes additional databases (included within OmniPath), which allow for a broader coverage of kinase–substrate interactions. However, users should be aware that larger kinase sets may include false positives, particularly for kinases with few mapped peptides. We therefore recommend interpreting individual kinase level results in conjunction with the kinase family level results and pathway enrichment output provided by the pipeline.

The 6-fold speedup over pamgeneAnalyzeR and 42-fold speedup over KRSA have practical implications beyond convenience. First, they enable immediate analysis at the point of data acquisition, supporting iterative experimental design where results from one experiment inform the next. Second, they support the creation of large labeled datasets suitable for training machine learning models for kinase activity prediction, a task that was previously impractical due to the computational cost of repeated full-pipeline analyses. More broadly, efficient large-scale kinase profiling may deepen our understanding of signaling networks and support kinase inhibitor development beyond oncology.

Several limitations should be noted. pyKinaXe currently supports only PamChip^®^ PTK and STK microarrays and is not directly applicable to other peptide microarray platforms. Also, both PTK and STK data need to be provided as input to pyKinaXe. Additionally, the statistical analysis, while robust, does not model dependencies between kinases that share substrate peptides, which may lead to correlated scores among closely related kinases. Furthermore, since phosphatase inhibitors are included within the sample preparation protocol for the PamChip^®^ microarray, the measured phosphorylation signals reflect only kinase-driven forward reactions. Phosphatase-mediated dephosphorylation (backward reaction) is therefore not reflected, which should be considered when interpreting pathway level results.

## Conclusion

We presented pyKinaXe, an open-source Python tool for one-click end-to-end kinase activity profiling from PamChip^®^ peptide microarray data. Our image analysis demonstrated robust and accurate microarray spot detection, which we benchmarked successfully against BioNavigator^®^. Validation on two independent datasets confirmed successful identification of most kinase families compared to KRSA, while achieving a 6-fold speedup over pamgeneAnalyzeR and 42-fold speedup over KRSA. Additionally, pyKinaXe allows the identification of individual kinases for higher resolution. pyKinaXe thus enables fast and reproducible kinase activity profiling, supporting both iterative experimental workflows and large-scale data generation for downstream pipelines.

## Data availability

All code required to reproduce the analyses presented in this paper is available at https://github.com/pykinaxe/pyKinaXe. The benchmarking dataset reanalyzed in this study is publicly available at Mendeley Data: Thiyagarajah et al. (2026), “Raw kinome array data - Differential impact of hepatitis delta virus replication and expression of viral antigens on the cellular kinome profile”, doi: 10.17632/ynp7f92n47.1.

## Key points

- Open-source tools for PamChip^®^ peptide microarrays have so far covered either image analysis or kinase inference. pyKinaXe combines both in one automated pipeline.
- On two public datasets, pyKinaXe’s spot intensities closely match those of the proprietary BioNavigator^®^ software, and its kinase family *z*-scores agree with those of KRSA computed from the same input.
- Beyond the kinase families reported by KRSA, pyKinaXe scores individual kinases using openly available kinase–substrate annotations. This layer is exploratory and best interpreted together with the family level results.
- pyKinaXe is open source and can also be used through a web interface.

## Author contributions statement

DW: Conceptualization, Methodology, Software, Validation, Formal analysis, Writing-Original Draft, Writing-Review, and Editing; EH: Conceptualization, Methodology, Validation, Writing-Review, and Editing; PK: Conceptualization, Methodology, Software, Validation, Writing-Original Draft, Writing-Review, Editing, Supervision, Project administration.

