## Supplementary Material for "pyKinaXe: a fast and robust turnkey kinase activity profiler"

### Validation of image analysis robustness

#### Robustness to image translation and rotation

The published source code for PamDx's gridding operator, 'pamsoft_grid', searches by default for rotations of the spot grid ranging from $-2°$ to $+2°$[4], indicating an estimation for the corresponding chip-insertion angular tolerance. We therefore assume that successful identification of all image spots within this range should cover all correctly manufactured and used PamChips. Our results show successful identification of all grid and reference spots, indicating the robustness of our image analysis. Example images of the applied transformations can be seen in Figure 1 for rotational transformation and in Figure 2 for translational transformation.

The code to generate these images can be found in the code repository under 'validate_papervalidate_robustness_of_image_analysis' and can be tested with own images if desired.

Additionally, we have tested image analysis on 13 different PTK-STK dataset pairs (>10000 images), with 100% success rate of accurate microarray spot identification, indicating statistical robustness of the image analysis pipeline [1].

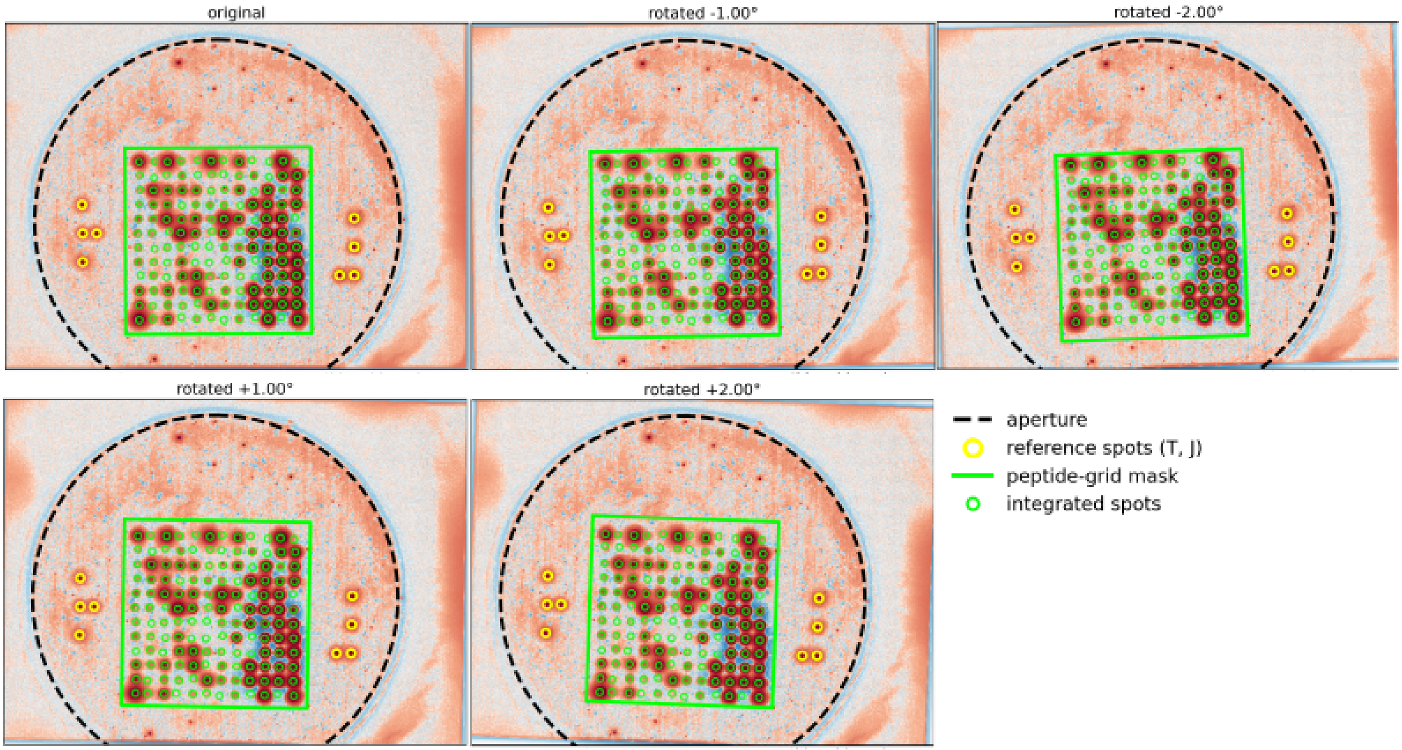


**Figure 1** Example images showing the successful identification of rotated images. The rotational transformation is applied between $-2^\circ$ and $+2^\circ$.

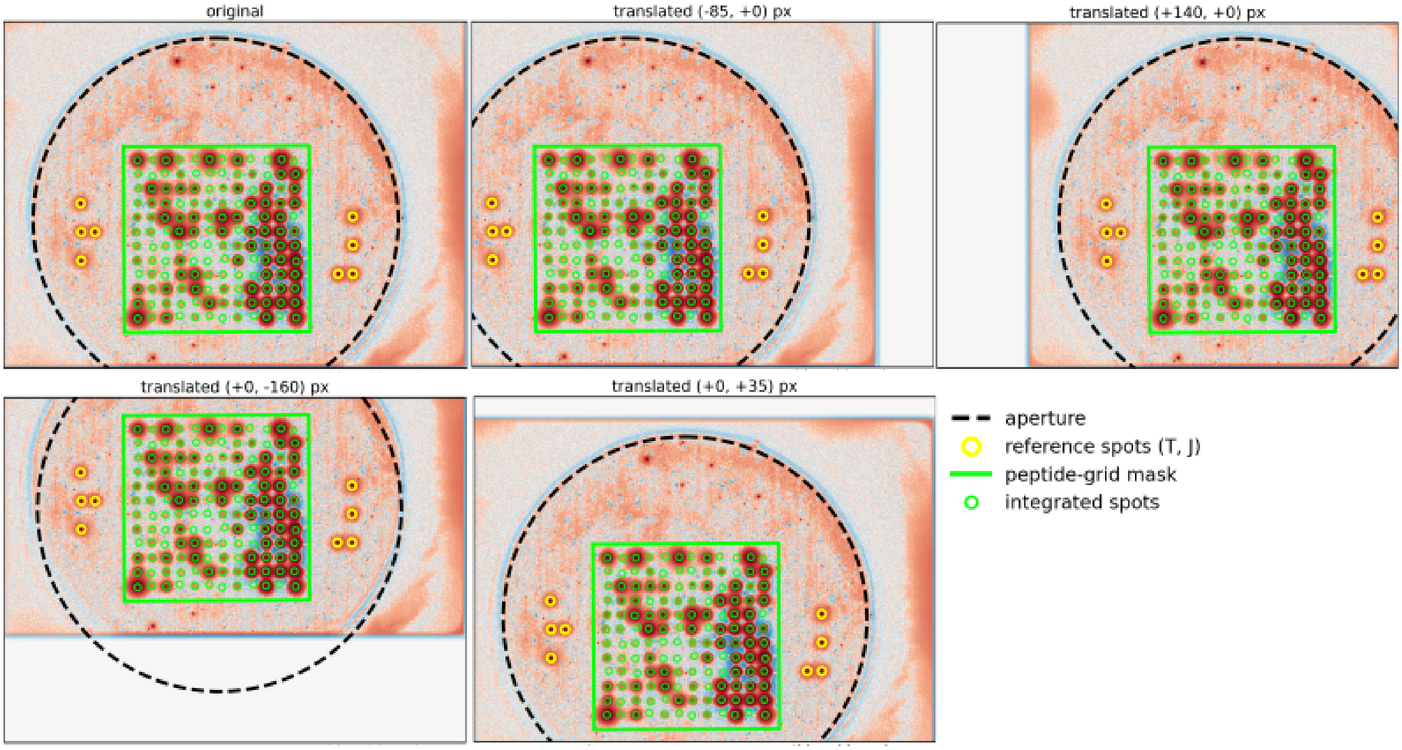


**Figure 2** Example images showing the successful identification of translated images. The translational transformation is applied until the grid of spots or the reference spots are too close to the border of the image.

### Validation of spot intensities against PamDx's BioNavigator

In pyKinaXe, there are two complementary approaches implemented for background (BG) estimation, the one based on morphological grey opening (MGO) and the other based on median BG estimated from rings around grid spots (Figure 3 for PTK, and Figure 4 for STK). The radius and width of each such ring were estimated to yield the best correlation with median intensities obtained via BioNavigator® (Figure 5).

The MGO-based approach estimates the base BG signal, but does not account for BG signal contributions (if any) from the neighboring spots (cross-talk). Such contributions become critical when saturated spots are bright enough to introduce additional BG signal to the neighborhood. In these cases, MGO-based BG estimation method underestimates the actual BG. The ring-based approach, on the other hand, yields a BG signal that reflects the actual intensity values in the raw data. In "cross-talk" cases described above, the ring-based approach therefore overestimates the actual BG.

The effects of the two BG estimation approaches can be understood from Figures 3 and 4.

The MGO-based BG estimate provides a baseline BG and is less affected by the neighboring spot signal, but can underestimate the true local BG in saturated-spot regions. A comparison of the median spot intensities (without MGO-based BG) with those estimated with BioNavigator® is shown in Figure 5a.

The ring-based approach estimates the local background from the rings surrounding each spot (orange in Figures 3 and 4). It is seen that in regions populated with very bright spots, signal from neighboring spots can extend into the rings, causing intensity cross-talk and leading to an overestimation of the background and consequently an underestimation of corrected spot intensities. This holds even in diagonal directions (Figures 3c and 4c) where the distance between neighboring spots is the largest. A comparison of the median intensities (without ring-based BG) with those obtained from BioNavigator® is shown in Figure 5b.

Thus, neither approach is expected to be universally optimal: the ring-based method tends to generally overestimate BG, whereas the MGO-based method tends to generally underestimate BG. As a compromise between these opposing biases and a rough approximation to actual BG, we, therefore, use the mean of the ring- and MGO-based BG estimates.

This averaged background estimation retained strong agreement with BioNavigator®-derived spot intensities (Figure 3 in the main text and Figure 5 c). Accordingly, all downstream validation analyses were performed using spot intensities corrected for this average BG estimation (default setting in pyKinaXe).

The dataset used for scatter plots in this section was published by Imami, A., & Vergis, J. (2026) and is publicly available at https://doi.org/10.5281/zenodo.20635299 [2].

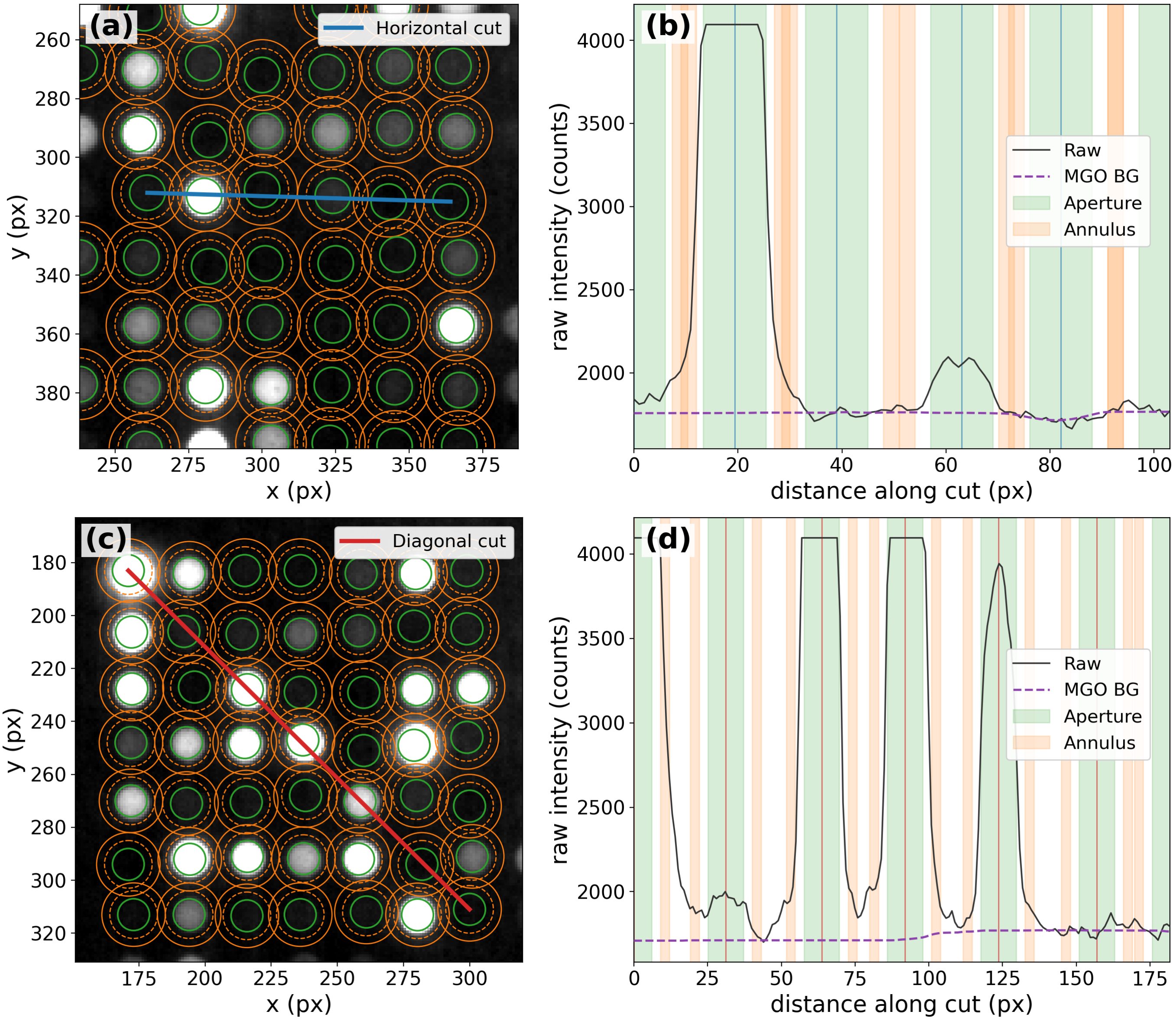


**Figure 3** Visualization of spot intensities on a PTK chip to estimate the background using the ring-based and MGO-based approach. (a) and (b) show plots for a horizontal cross-section, (c) and (d) for a diagonal cross-section.

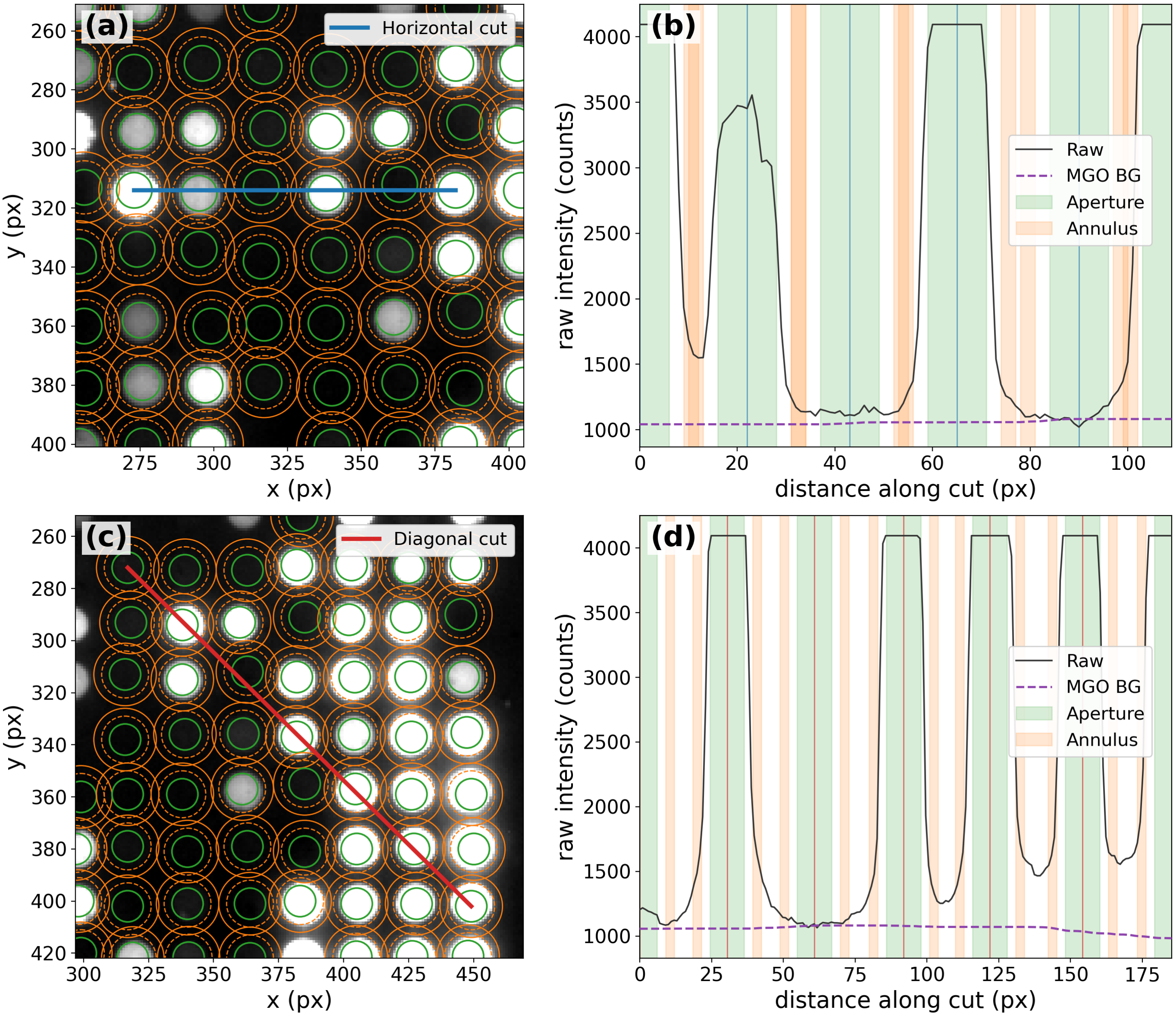


**Figure 4** Visualization of spot intensities on a STK chip to estimate the background using the ring-based and MGO-based approach. (a) and (b) show plots for a horizontal cross-section, (c) and (d) for a diagonal cross-section.

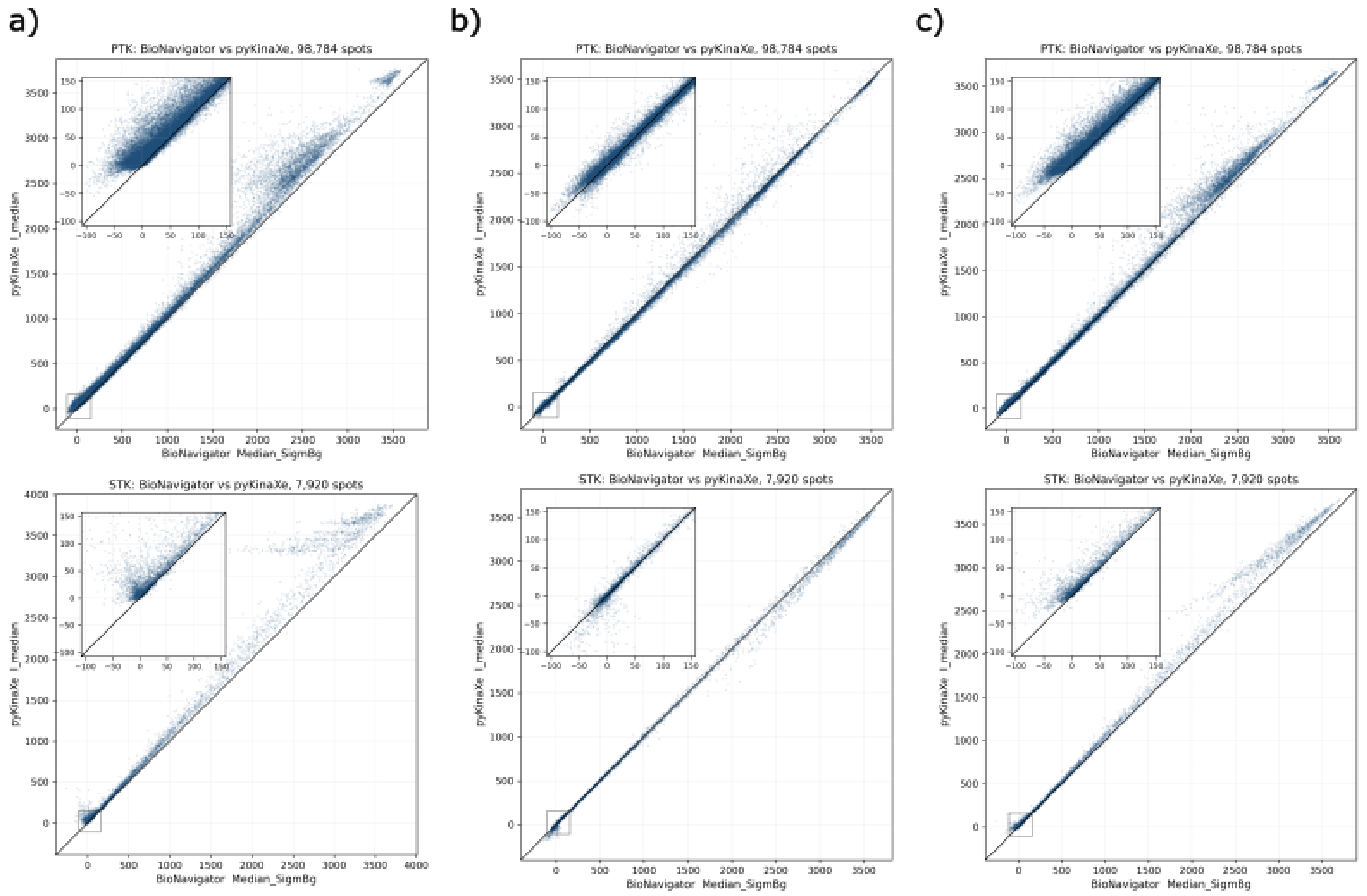


**Figure 5** Scatter plots comparing spot intensities produced by pyKinaXe (y-axis) against spot intensities produced by BioNavigator® (x-axis). (a) shows spot intensities with background estimation using the MGO-based approach. (b) shows intensities estimated using the ring-based approach. (c) shows the plot for the averaged estimation.

### Validation of UKA on a second dataset

We validated pyKinaXe's UKA against the second dataset by Imami et al. [2].

In the corresponding paper Lee et al. [3] investigated how acute voluntary wheel running alters kinase signaling in the rat brain, focusing on the dorsal striatum and hippocampus, regions involved in movement, motivation, and exercise-responsive plasticity. The authors profiled kinase activity using PamDx PamStation12 STK/PTK kinome arrays, performed kinase anylsis with KRSA, UKA, and KEA3, and integrated these outputs with Creedenzymatic plus downstream pathway and protein-network analyses. The study found region-specific signaling shifts, including decreased serine/threonine kinase activity, increased tyrosine kinase activity, altered IKK/PKD and CSK/EGFR/VEGFR-related pathways, and NF-kB changes linked to running distance, suggesting that acute exercise rapidly engages brain kinase networks tied to exercise behavior. The paper is publicly available at https://doi.org/10.1371/journal.pone.0321596.

As shown in the results section of our paper, we validate the image analysis output of pyKinaXe against the image analysis output of BioNavigator®. Therefore, we compare the spot intensities produced by the image processing analysis against the output of BioNavigator®'s image analysis (Figure 6). The results show both for PTK (a) and STK (b) a similar behavior of spot intensities compared to the results from BioNavigator®.

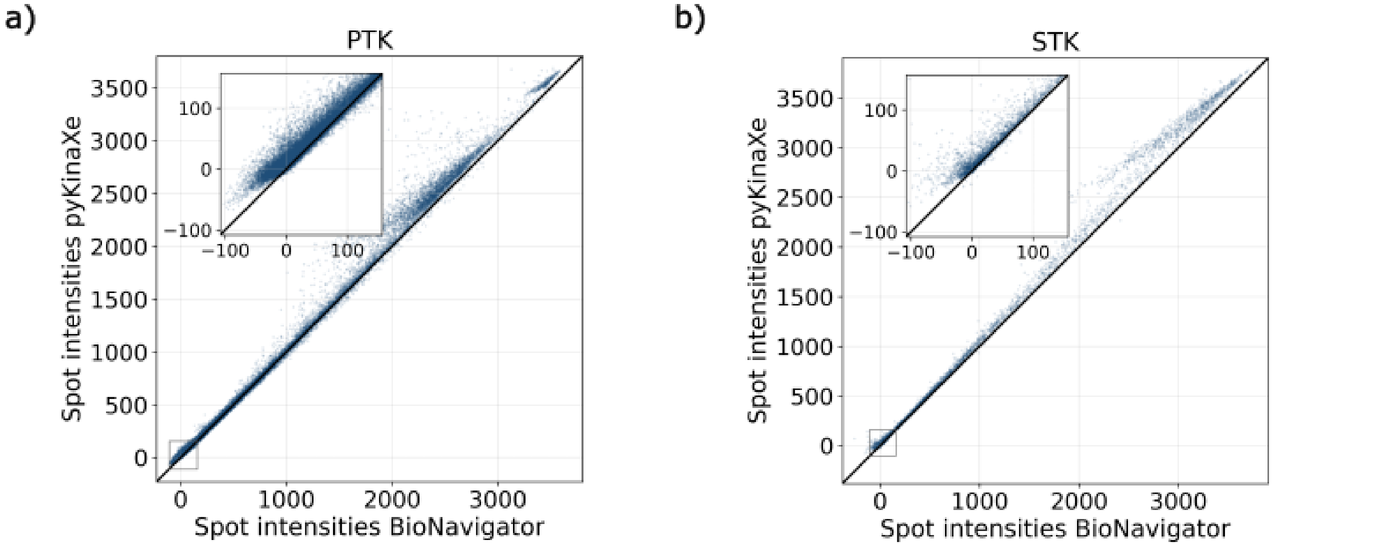


**Figure 6** Comparison of spot intensities following image analysis between the output of pyKinaXe (y-axis) and BioNavigator® (x-axis). A magnified view shows the spot intensities at lower intensities. (a) shows the spot intensities for PTK chips, and (b) shows the intensities for STK chips.

Additionally, we performed the upstream kinase analysis for kinase families both using pyKinaXe and KRSA (on pyKinaXe image analysis input, similar to the validation in the paper). The results of this comparison are shown in Figure 7.

#### Making the second dataset work on pyKinaXe

To use the second validation dataset with pyKinaXte, the end user needs to make certain changes and corrections to the data:

- As the authors mention in the paper, not all generated data can be used for the analysis. Due to failure of a chip, only two out of the three technical replicates for the STK chips were used for analysis [3]. Therefore, images from STK chip number 710230320 need to be removed from the data.
- Spelling errors in the sample annotation file need to be corrected to make sure all samples have matching names.
- Additional changes in the sample name need to be performed, in order to tell pyKinaXe which sample is control (c) or test (t).
- Correcting artifacts in the array layout files. Peptides with an inserted whitespace, like 'JAK1_ 1027_1039' or 'MK07_ 212_224' need to be adjusted.

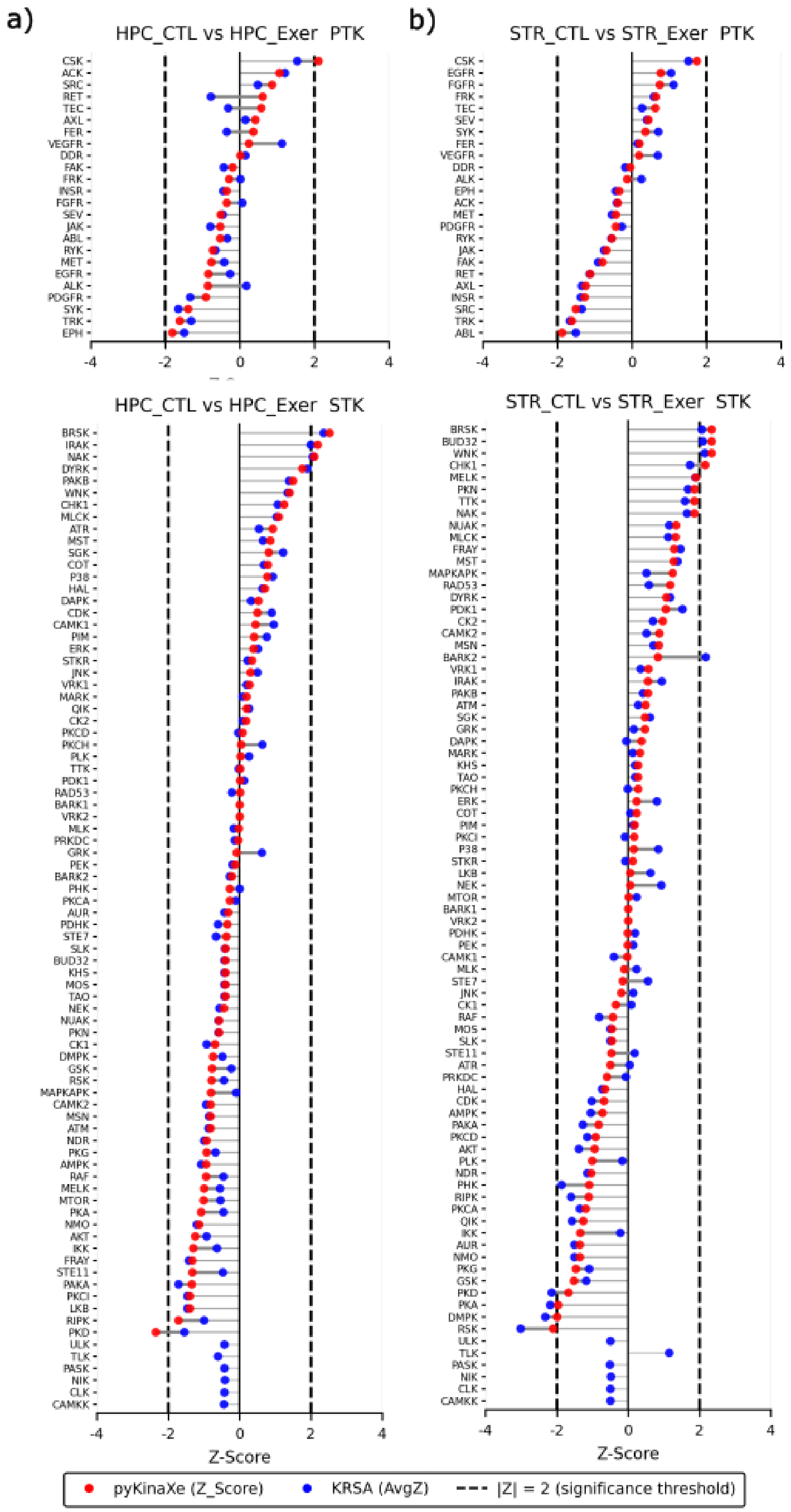


**Figure 7** Comparison of the $z$-scores of kinase families between pyKinaXe and KRSA based on the pyKinaXe image analysis output. The x-axis shows the $z$-score, and the y-axis shows the various kinase families (sorted by z-score). (a) shows waterfall plots comparing HPC control to HPC test for both PTK and STK chips. (b) shows the results for STR control to STR test. The blue points represent the $z$-scores from KRSA, and the red points represent the $z$-scores from pyKinaXe. The significance levels (dashed vertical lines) are $-2$ and $+2$.
